# Memory-based incremental parameter updating of a generic stochastic plant epidemic model

**DOI:** 10.1101/2025.07.11.664325

**Authors:** Israël Tankam Chedjou

**Affiliations:** L’Institut Agro Rennes-Angers

**Keywords:** *Stochastic epidemic modelling*, *Plant disease surveillance*, *Real-time parameter estimation*, *Particle filtering*, *Memory-based sampling*

## Abstract

In plant-disease surveillance, timely and accurate estimation of transmission parameters is critical for informed decision-making. Here, I present a sequential Monte Carlo method that incrementally updates key parameters of a stochastic compartmental epidemic model as new incidence data are collected daily. My approach couples a Gillespie stochastic simulation algorithm for disease epidemiology with a memory-based particle-resampling scheme that allows real-time inference of primary and secondary infection rates even when observed infection counts are low. Using a synthetic outbreak to mimic typical field epidemics, I show that posterior means for transmission parameters converge to within an average of *∼* 10% of true values by Days 10-15 post-introduction. Concurrently, ensemble-based short-term forecasts achieve *R*^2^ *∼* 0.8 by Day 2 and exceed *R*^2^ *≈* 0.93 by Day 30. Computational costs remain modest; each daily update completes in under 4 seconds on standard hardware, highlighting the feasibility of integrating this method into automated surveillance platforms. While validation against synthetic data shows strong performance, I discuss potential challenges in real-world applications, including real data, model misspecification, latent infection dynamics, and spatial heterogeneity. This sequential-Bayesian approach provides a scalable, uncertainty-aware solution for real-time parameter estimation and forecasting in stochastic plant-epidemic systems, laying the groundwork for adaptive management of crop-disease outbreaks.

## 1 Introduction

Mathematical models have become indispensable tools in epidemiological decision-making, particularly in the management of plant diseases where timely and informed interventions are essential to preventing or limiting outbreaks. Their value lies in their ability to synthesize complex biological processes and forecast disease dynamics under various scenarios. To guide effective interventions, epidemiological models must excel in two areas. First, they need to quantify uncertainty: every input, as well as demographic stochasticity [12], carries variability, and nonlinear interactions can amplify small errors. Stochastic formulations (e.g., Gillespie compartmental models [15] or stochastic differential equations) produce confidence envelopes around incidence curves, giving decision-makers clear margins of error [25, 13]. Second, models rely on rate parameters that set epidemic speed and scale, and they need to be excellently estimated. While “speed” rates (e.g. latent-to-infectious transitions or mortality) can be estimated from cohort time data (e.g., the reciprocal of mean incubation), transmission parameters (like contact rate, infection probability per encounter, acquisition and inoculation efficiencies) are far harder to observe [24]. Consequently, these infection rates are often only roughly approximated or assumed instantaneous in worst-case scenarios [26]. Yet small errors in transmission rates can yield large mispredictions of outbreak timing and size, undermining model-based decisions.

Donnelly and Gilligan [7]and Donnelly et al. [8] introduced an innovative approach for estimating infection rates using “access-period” laboratory assays for vector-borne plant viruses. While this technique is readily extensible to a broad spectrum of plant pathogens, it remains fundamentally dependent on pre-outbreak laboratory data, a resource that is often unavailable when an epidemic first emerges. Moreover, even when such assays are conducted, their accuracy can be compromised by the inherent mismatch between controlled experimental settings and complex field environments. Factors such as climatic or thermal stress on hosts and vectors, as well as genetic and varietal diversity not captured in laboratory cohorts, can lead to parameter estimates that do not faithfully represent in-situ transmission dynamics [4]. A posteriori outbreak estimation methods address these challenges by inferring epidemic parameters from observed field dynamics after an outbreak has begun. As Thompson et al. [27] emphasized, dedicating time to real-time data collection during the early stages of an epidemic is crucial for informing the most effective response.

Yet, most existing parameter inference techniques for dynamical systems, including least-squares, are designed for batch-based calibration, assuming full access to the complete time series of observations. In the context of plant epidemiology, the standard practice remains the retrospective fitting of deterministic or stochastic models to full disease progress curves using nonlinear least squares or maximum likelihood estimation. While Bayesian methods such as Sequential Monte Carlo (SMC) [9] or information-theoretic approaches like Information Field Theory (IFT) [11] can be applied for sequential parameter estimation, where estimates are refined dynamically at each new observation, they can be computationally heavy. Computationally light sequential parameter updates have received very limited attention, especially in plant disease modelling. Similarly, while machine learning methods such as LSTM [1] can help with sequential updating of disease trajectories, they focus on forecasting rather than interpretable parameter inference [29, 1] and may still be computationally heavy.

This study describes a generic Gillespie model for plant disease epidemiology and a memory-based particle resampling scheme to enable efficient, real-time updating of transmission parameters. The proposed approach combines the stochastic simulation algorithm with a sequential Monte Carlo framework that uses the posterior distribution from the previous day to generate proposals for the next, thereby concentrating computational effort in high-posterior regions and accelerating convergence. I validate the method using synthetic data generated from a typical plant disease outbreak, demonstrating rapid and accurate parameter estimation along with reliable short-term forecasts. The paper is structured as follows: Section 2 details the epidemic model and the sequential parameter update algorithm. Section 3 presents the results of the synthetic validation, including forecast accuracy, parameter convergence, and computational performance. Section 4 discusses the implications, limitations, and potential extensions of the work.

## 2 Material and methods

### 2.1 The standard model

I state that the following equation (1) proposed equation captures the essential dynamics of plant disease epidemics while remaining parsimonious:

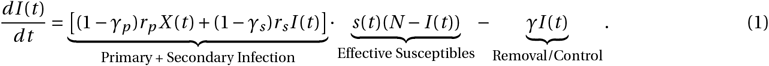

Where, *I* (*t*) is the infected plant population, *N* is the total host population in the field, (*N ™ I* (*t*)) is the susceptible plant population, *X* (*t*) is the density of initial inoculum, *r*_*p*_ and *r*_*s*_ are the rates of primary and secondary infection respectively, *s*(*t*) is a measure of the change in susceptibility of the host, *γ*_*p*_ and *γ*_*r*_ are measures of control efforts, and *γ* represents the death and/or the removal of infected individuals from the host population.

This equation encapsulates the essential dynamics of plant disease epidemics by integrating four fundamental processes: primary infection, secondary infection, host susceptibility, and removal or control measures. Below, I justify its universality and address potential extensions.

#### Universality of the core processes

This equation integrates four fundamental processes: primary infection, secondary infection, host susceptibility, and removal or control measures.

1. Primary Infection The term (1 – *γ*_*p*_)*r*_*p*_ *X* (*t*) represents the rate of primary infection, where *X* (*t*) denotes the external inoculum source, such as soilborne pathogens or airborne spores. This component initiates the epidemic by introducing the pathogen into the host population. The factor (1 *™ γ*_*p*_) accounts for the effectiveness of control measures targeting primary infection sources, including fungicide applications, crop rotation, or sanitation practices. This modelling approach aligns with the SIRX framework, which incorporates both primary and secondary infection sources to simulate plant disease dynamics effectively [16, 6].
2. Secondary Infection The term (1 –*γ*_*s*_)*r*_*s*_ *I* (*t*) captures the rate of secondary infection, where *I* (*t*) is the number of infected individuals contributing to the spread of the disease within the host population. Secondary infection is characteristic of polycyclic diseases, where multiple infection cycles occur within a single growing season. Control measures targeting secondary infection, represented by *γ*_*s*_, include practices such as pruning, or the use of resistant cultivars. This component is crucial for modeling the progression of diseases that exhibit density-dependent transmission dynamics [20].
3. Host Susceptibility The factor *s*(*t*)(*N* – *I* (*t*)) represents the effective number of susceptible hosts, where *N* is the total host population and *s*(*t*) is a time-dependent function reflecting variations in host susceptibility due to factors like age, environmental conditions, or phenological stages. This term allows the model to account for changes in susceptibility over time, such as increased resistance in mature plants or heightened vulnerability during specific growth stages [22]. Incorporating host susceptibility dynamics enhances the model’s ability to simulate real-world disease progression accurately.
4. Removal or Control Measures The term –*γI* (*t*) represents the removal or control of infected individuals from the host population. This encompasses various management strategies, including the physical removal of infected plants, application of systemic treatments, or natural death of hosts. The parameter *γ* quantifies the rate at which infected individuals are removed, thereby reducing the overall disease burden.

#### Why “biggest” models can be seen as complexifications

The generalized plant disease epidemic model (1) offers a streamlined yet comprehensive framework for understanding disease dynamics. While various extensions introduce additional complexities, many of these can be effectively approximated within the existing model structure. Below, I examine some common model extensions and discuss how their effects can be captured without significantly increasing model complexity.

1. 1. Latent periods (SEI Models) Extensions like the SEI (Susceptible-Exposed-Infectious) model introduce an “exposed” compartment to represent the latent period between infection and infectiousness. While this adds biological realism, especially for diseases with significant incubation periods, the generalized model can approximate latency by adjusting parameters such as the primary infection rate (*r*_*p*_), secondary infection rate (*r*_*s*_), or the susceptibility function (*s*(*t*)). These adjustments can delay the onset of infectiousness, effectively capturing the essence of a latent period without adding a separate compartment.
2. Spatial heterogeneity Incorporating spatial heterogeneity often involves complex models using partial differential equations or network-based approaches to simulate disease spread across landscapes. However, spatial effects can be implicitly represented in the generalized model by modifying parameters like the external inoculum source (*X* (*t*)) or the secondary infection rate (*r*_*s*_) to reflect spatial gradients or contact rates. This approach captures the influence of spatial factors on disease dynamics without the need for spatially explicit modelling. While the model may become less precise, the count of infected hosts might not be too much influenced if the infection rates *r*_*s*_ and *r*_*p*_ are well calibrated; which is the purpose of this paper.
3. Host demography Models that incorporate host demography add birth and death terms to account for changes in the host population over time. While important for long-term or perennial systems, these factors may be negligible in short-term epidemics where the host population remains relatively constant. In such cases, the generalised model can assume a fixed host population (*N*), simplifying the analysis without significantly compromising accuracy.
4. Stochasticity Introducing stochastic elements into disease models accounts for random variations and uncertainties in disease spread. As argued in the introduction, I value the stochasticity, and I introduce the stochastic form of model (1) through the Gillespie algorithm [15] in Table 1.

**Table 1.** Gillespie algorithm form of the standard epidemics model (1). reaction channels, propensities, and state changes. Modelling details are explicited in Appendix A and the forms of *X* (*t*) and *s*(*t*) are explicited in equations (3) and (2) respectively.

| Reaction $j$ | Propensity $a_j(I, t)$ | Change $\Delta I_j$ |
| --- | --- | --- |
| 1. Infection | $[(1 - \gamma_p)r_p X(t) + (1 - \gamma_s)r_s I] s(t)(N - I)$ | +1 |
| 2. Removal | $\gamma I$ | -1 |

I assume without loss of generality that control rates *γ*_*p*_ and *γ*_*s*_ are equal to 0. As often in the literature [22], I assume that *s*(*t*) takes the form:

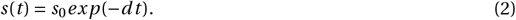

I also write:

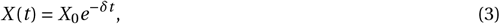

The form (3) is derived from the general form 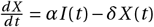, by setting the production rate *α =* 0 during the growing season. This simplification is biologically justified: primary inoculum (e.g., overwintering spores, soilborne pathogens, or crop residue-borne pests and pathogens) originates from pre-season sources and degrades at rate *δ*, while new infections eventually arise mainly from secondary inoculum produced by within-season infected plants (*I* (*t*)). By neglecting external inoculum replenishment (*α =* 0), I adhere to the ecological principle that primary and secondary infection phases are temporally distinct in plant pathology, a framework validated in several systems [5, 28].

### 2.2 Parameter and data

This section describes the parameter values and synthetic data used to validate the incremental parameter estimation method for the plant disease epidemic model (Table 1). All parameters are grounded in realistic plant pathology scenarios, and synthetic data provides a controlled environment for proof-of-concept testing.

#### Model Parameters

The epidemic model is governed by the following fixed parameters:

- Total host population: *N =* 100 (finite population of susceptible plants)
- Initial inoculum: *X*_0_ *=* 25.0 (units of pathogen propagules)
- Inoculum decay rate: *δ =* 0.03 (exponential decay of primary inoculum)
- Removal/recovery rate: *γ =* 0.1 (implies average infectious period of 1/*γ =* 10 days)
- Initial infected hosts: *I*_0_ *=* 1 (single introductory infection)
- Host susceptibility decay: *s*(*t*) *= e*^−0.001*t*^ (slow reduction in host susceptibility over time)

#### Synthetic Data Generation

The time-series of infected hosts *I* (*t*) was generated by numerically solving the deterministic model:

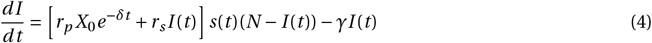

using a 4th-order Runge-Kutta solver with Δ*t =* 0.1 days. In addition to the parameter values above, I consider *r*_*p*_ *=* 8.5*×*10^−4^ and *r*_*s*_ *=* 1.1*×*10^−3^ to generate the synthetic infected counts (Table 2). The solution was sampled at daily intervals *t =* {0, 1,…, 30}, rounded to integers (reflecting discrete host counts), and perturbed with Gaussian noise *𝒩* (0, 0.1) to mimic observation error for example.

**Table 2.** Synthetic time-series of infected hosts *I* (*t*)

|  |  |  |  |  |  |  |  |  |  |  |  |  |  |  |  |  |
| --- | --- | --- | --- | --- | --- | --- | --- | --- | --- | --- | --- | --- | --- | --- | --- | --- |
| $t$ (days) | 0 | 1 | 2 | 3 | 4 | 5 | 6 | 7 | 8 | 9 | 10 | 11 | 12 | 13 | 14 | 15 |
| $I(t)$ | 1 | 3 | 5 | 8 | 9 | 10 | 13 | 14 | 14 | 16 | 17 | 18 | 19 | 19 | 20 | 21 |
| $t$ (days) | 16 | 17 | 18 | 19 | 20 | 21 | 22 | 23 | 24 | 25 | 26 | 27 | 28 | 29 | 30 | |
| $I(t)$ | 22 | 23 | 23 | 23 | 25 | 25 | 25 | 25 | 25 | 26 | 26 | 26 | 26 | 26 | 26 | |

Synthetic data enables a controlled validation; indeed, known ground-truth parameters (*r*_*p*_, *r*_*s*_) allow rigorous evaluation of estimation accuracy.

#### Incremental Data Protocol

For methodology testing, *I* (*t*) is delivered sequentially:

- At *t*_*k*_ *= k* days, only {*I* (0), *I* (1),…, *I* (*k*)} is available
- Parameters *r*_*p*_ and *r*_*s*_ are re-estimated upon receipt of each new datum (Section 2.3)
- Performance metrics are evaluated at each *t*_*k*_

The synthetic time-series provides a reproducible benchmark for evaluation the incremental parameter update method.

### 2.3 Parameter update and prediction

This section details the mathematical framework and implementation of the sequential Monte Carlo (particle-filtering) estimator for the transmission parameters *r*_*p*_ (primary inoculum rate) and *r*_*s*_ (secondary transmission rate) in the stochastic model Tables 1 with time-varying inoculum *X* (*t*) and susceptibility *s*(*t*). The model process evolves as a continuous-time Markov jump process with two types of transitions:

- Infection: *I → I +* 1 at instantaneous rate

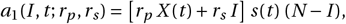
- Removal: *I → I ™* 1 at instantaneous rate

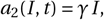

#### Daily observation

I assumed discrete-time surveillance: at each integer day *k =* 1, 2,.. ., I observe the count of infected hosts *Y*_*k*_. Denote by 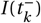 the value of *I* (*t*) just prior to day *k* (i.e. immediately before the *k*-th observation). I model

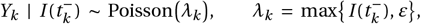

where *ε >* 0 is a small regularization constant to avoid zero-mean degeneracy.

#### Sequential Bayesian inference via Particle Filtering

Because data arrive incrementally (day-by-day), I adopt a sequential Monte Carlo (SMC) approach specifically, a bootstrap particle filter to approximate the Bayesian posterior

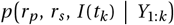

in real time [18, 10, 3]. Crucially, I incorporate a memory-based sampling step so that after assimilating data on day *k*, the particle set used on day *k +* 1 is drawn from a Gaussian approximation to the day-*k* posterior. This accelerates convergence and concentrates computation in high-posterior regions.

#### Particle representation

At the start of each day *k*, I maintain *N*_*p*_ particles

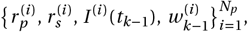

where 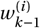 approximates the posterior weight

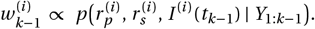

The static parameters (*r*_*p*_, *r*_*s*_) are treated as unknowns with prior support in **ℝ**^*+*^ *×***ℝ**^*+*^, and the dynamic state is *I* ^(*i*)^(*t*_*k™*1_).

#### Propagation (prediction)

Given particle *i* at day *k ™* 1, I have

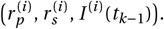

To predict the state at time *t*_*k*_, I simulate the continuous-time Markov jump process from *t*_*k™*1_ to *t*_*k*_ via the Gillespie algorithm [14]. Let

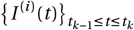

be the resulting trajectory. I then set the propagated latent state just before observation *k* as

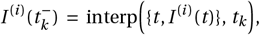

where interp denotes linear interpolation over the jump times to evaluate *I* (*t*) exactly at *t*_*k*_.

#### Incremental weight update

Upon observing *Y*_*k*_ *= y*_*k*_, I update each particle’s weight by multiplying its prior weight by the Poisson likelihood:

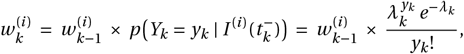

With

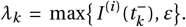

I then normalize all weights so that 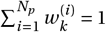

#### Resampling and memory-based sampling

To prevent weight degeneracy, I compute the effective sample size (ESS):

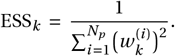

If ESS_*k*_ *< τ N*_*p*_ (e.g. *τ =* 0.3), I perform systematic resampling [21] to produce an unweighted set 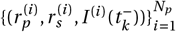 each with new weight 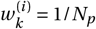

However, instead of carrying these particles forward directly, I introduce a memory-based sampling step that uses the day-*k* posterior distribution to generate new proposals for day *k +* 1. Concretely:

1. 1. Compute the posterior mean of (*r*_*p*_, *r*_*s*_):

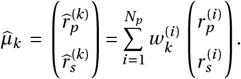
2. Compute the weighted covariance matrix of the particles around 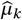:

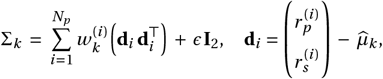

with a small inflation term *ϵ ≪* 1 (e.g. 10^*™*12^) to ensure positive definiteness.
3. Draw a new set of *N*_*p*_ proposals for (*r*_*p*_, *r*_*s*_) at day *k +* 1 from a multivariate normal distribution centered at 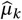 with covariance Σ_*k*_ :

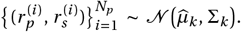

I then enforce positivity by taking 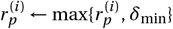 and likewise for 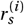, with *δ*_min_ *=* 10^*™*8^.
4. Reset all weights to 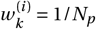 The latent states 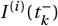 are carried forward to serve as the starting point for propagation on day *k +* 1.

This memory-based resampling ensures that on day *k +* 1, particles are already clustered near the high-posterior region identified on day *k*, which accelerates convergence and improves sampling precision.

#### Parameter estimation

At the conclusion of the last survey day *k*, the SMC algorithm yields the Posterior mean estimates:

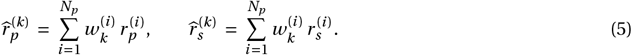

Because each new observation *Y*_*k*_ is assimilated immediately and the particle set is re-centered around the previous day’s posterior via memory-based sampling, this algorithm provides real-time refinement of (*r*_*p*_, *r*_*s*_)and tight predictive distributions with minimal wasted computation in low-posterior regions. Box 2 presents a least-squares parameter update method that performs well for these model and the data, but lacks generality.

##### Box 1

**Summary of the sequential algorithm**

At each day *k =* 1, 2,.. .:

1. **Propagation:** For each particle *i*, simulate *I* ^(*i*)^(*t*) from *t*_*k™*1_ to *t*_*k*_ via Gillespie’s algorithm and set 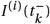 by interpolation.

2. **Weight Update:** Observe *Y*_*k*_ *= y*_*k*_. Update weights

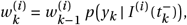

then normalize.

3. **Resampling Decision:** Compute ESS_*k*_. If ESS_*k*_ *< τ N*_*p*_, perform systematic resampling to obtain unweighted particles.

4. **Memory-Based Sampling:** Compute the weighted mean 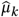 and covariance Σ_*k*_of 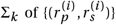 Draw new 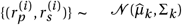, enforce positivity, and reset weights to uniform. Carry forward the latent counts 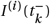.

5. **Parameter Estimation:** Record the posterior means

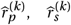

as the day-*k* estimates.

6. **Predictive Ensemble:** Generate *M* independent Gillespie simulations using 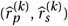. Compute 2.5%-97.5% quantiles across these *M* trajectories for uncertainty envelopes.

7. **Fit Diagnostics:** For the first *k* days, compare the predictive mean at times *t =* 0, 1, 2,…, *k* to observations *Y*_1:*k*_, computing:

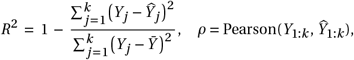

along with the associated *p*-value for testing zero correlation [3].

##### Box 2

**Least-squares parameter update**

With this simple generic model (Table 1), I can estimate *r*_*p*_ (primary infection rate) and *r*_*s*_ (secondary infection rate) to field surveillance data via a least-squares approximation of the underlying mean-field ordinary differential equation (ODE) (1). By simulating the ODE forward with a given (*r*_*p*_, *r*_*s*_), I obtain a smooth trajectory *î*(*t*) and define the sum of squared errors

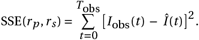

Minimizing

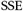

yields maximum-likelihood estimates under a Gaussian-error approximation.

**When does it perform well for this model**

With a moderate to high host population (*N ≥* 100), stochastic Gillespie realizations cluster tightly around their mean. The ODE thus captures the expected behaviour accurately, making residuals roughly homoscedastic and Gaussian. Additionnaly, the deterministic trajectory *Î* (*t*) depends smoothly on (*r*_*p*_, *r*_*s*_). This yields an SSE surface with a single, well-defined minimum, enabling rapid convergence. Even more since evaluating the ODE is orders of magnitude faster than simulating thousands of stochastic paths in an inner optimization loop. But this good performance is realized upon approximate linearization, which leads to the escape from the non-linearity limitation I mention further. In detail, the number of infected hosts might be low for most of the observation window. Hence, there is an early-stage dominance of the linear regime. Indeed, when *I* is small relative to *N* in the early stages of the epidemics,

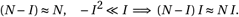

Thus the quadratic “saturation” term *™r*_*s*_ *s*(*t*) *I* ^2^ contributes only a small correction to the overall infection rate. In effect, the system behaves almost like a linear birth-death process over the range of your data, where the mean-field ODE is an excellent proxy for **𝔼** [*I* (*t*)].

**Limitations and lack of generality**

Despite its success here, the mean-field least-squares approach relies on two critical assumptions that often fail in more complex epidemic or ecological models. Indeed, for many processes (e.g. spatially explicit spread, age-structured populations, or models with higher-order interactions) the exact mean ODE is unknown or intractable. Additionally, in models with strong non-linearities,

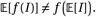

The average of stochastic trajectories does not follow the deterministic solution, leading to biased parameter estimates if one fits the ODE directly. A classic example is a model with infection term *∝ I* ^2^, for which **E**[*I* ^2^] *=* Var(*I*) *+* (**E**[*I*])^2^, so the deterministic ODE underestimates the true infection pressure.

**Outlook**

To take advantage of the speed of the least-square approximation when the mean-field equation is a good proxy of the stochastic model, I can employ a two-stage inference: using the mean-field fit as a warm start, when the epidemic level is low (e.g *I* /*N <* 0.3), then change it for a more rigorous parameter update method. Such extension retains the conceptual appeal of least-squares fitting while overcoming the restrictive assumptions inherent to the classic mean-field approach.

#### 2.4 Forecasting and goodness-of-fit

By simulating *M* independent Gillespie trajectories from time *t*_*k*_ onward using the estimated 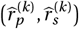 in eq (5),I obtain a posterior predictive distribution on future infected counts; 2.5% and 97.5% quantiles form a 95% credible envelope.

The coefficient of determination *R*^2^ and Pearson correlation *ρ* (with *p*-value) compare model mean trajectory predictions to the cumulative observations *Y*_1:*k*_. These metrics are particularly appealing as fitness metrics in this context because they quantify, in a simple and interpretable manner, how closely the model’s predicted mean trajectory matches the observed infection counts: *R*^2^ explicitly measures the proportion of variability in the observed data explained by the predictive mean [19], while Pearson’s *r* assesses the strength of linear association between predicted and observed values [23]. Both statistics are readily computed at each sequential update and provide immediate diagnostics of model calibration (i.e., whether the model is systematically under-predicting) and precision (i.e., how tightly the predictions track the day-to-day changes). Because my primary interest is in tracking the trajectory of cumulative infected hosts (continuous) these metrics capture the essential goodness-of-fit without requiring more complex scoring rules. Possible alternatives include point-wise error measures such as root-mean-square error (RMSE) or mean absolute error (MAE), which directly quantify average forecast error in the original count scale but do not summarize explained variance [30]; likelihood-based criteria (e.g., log-likelihood, AIC, BIC) [2]; and fully probabilistic scoring rules such as the continuous ranked probability score (CRPS) or the Dawid-Sebastiani score, which evaluate the entire predictive distribution rather than just its mean [17].

## 3 Results

To infer the infection and secondary transmission rates over time, I implemented a particle filtering algorithm tailored to the stochastic dynamics of the epidemic process. The filter was initialized with 200 particles, each representing a pair of candidate parameters (*r*_*p*_, *r*_*s*_) sampled from a uniform prior. At each daily observation step, 10 independent Gillespie simulations were run per particle to estimate the likelihood of the observed number of infections, assuming a Poisson observation model. The filter employed systematic resampling whenever the effective sample size (ESS) dropped below 30% of the particle count, ensuring robustness against particle degeneracy.

After resampling, particles were re-initialized from a multivariate normal distribution centered on the posterior mean with a covariance estimated from the weighted particle ensemble. This memory-based propagation allows the filter to adapt dynamically to observed data while maintaining sufficient variability in the parameter space. To evaluate model performance and parameter convergence, I recorded the posterior means, 95% credible intervals, and goodness-of-fit metrics (R^2^ and Pearson correlation) at each assimilation step. At each step, I also plot the forecast trajectory upon 100 simulations of the Gillepsie algorithm with mean parameter estimates.

### 3.1 Real-time forecasts

The memory-based particle filter rapidly learns the underlying dynamics.

1. **Day 1 (Fig. 1A)**. With a single noisy observation *Y*_1_ *=* 1, the posterior remains near the uniform prior mean in [10^*™*5^, 10^*™*3^]. The ensemble mean peaks around 10 infected hosts at *t ≈* 15 days, and the 95 % envelope spans roughly [0, 18] hosts for 0 *≤ t ≤* 100. The lone data point at (*t =* 0, *Y*_1_ *=* 1) lies at the lower edge of this envelope.
2. **Day 5 (Fig. 1B)**. After assimilating {1, 3, 5, 8, 9}, the posterior-mean parameters shift to 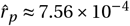 and 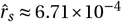 (cf. Fig. 3). The ensemble mean now peaks near 17-18 hosts at *t ≈* 22, and the 95 % envelope peak range *≈* [8, 30]. All five observed points lie within this envelope.
3. **Day 10 (Fig. 1C)**. With ten observations {1, 3, 5, 8, 9, 10, 13, 14, 14, 16}, 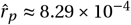 and 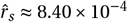 The predictive mean peaks around 20-21 hosts, and the envelope around *≈* [10, 32]. All ten blue points lie close to or within *±*1 host of the red mean, indicating excellent calibration.
4. **Day 15 (Fig. 1D)**. By Day 15 (*Y*_15_ *=* 20), 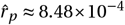 and 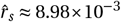, approaching the true generating values. The 95 % envelope at *t ≈* 15 spans [9, 38], and all fifteen observations remain inside, and close to the red mean.
5. **Day 20 (Fig. 1E)**. After twenty data points (*Y*_20_ *=* 23), 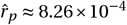 and 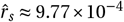. All twenty counts lie centrally within the band, demonstrating robust performance despite observation noise.
6. **Day 30 (Fig. 1F)**. At the end of the 30 days (*Y*_30_ *=* 26), 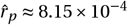 and 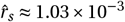, close to the true values 8.5 *×* 10^*™*4^ and 1.1 *×* 10^*™*3^ from the mean-field generator (2.2). Nearly all 30 observations lie within *±*1 host of the mean, indicating high confidence.

**Figure 1.**
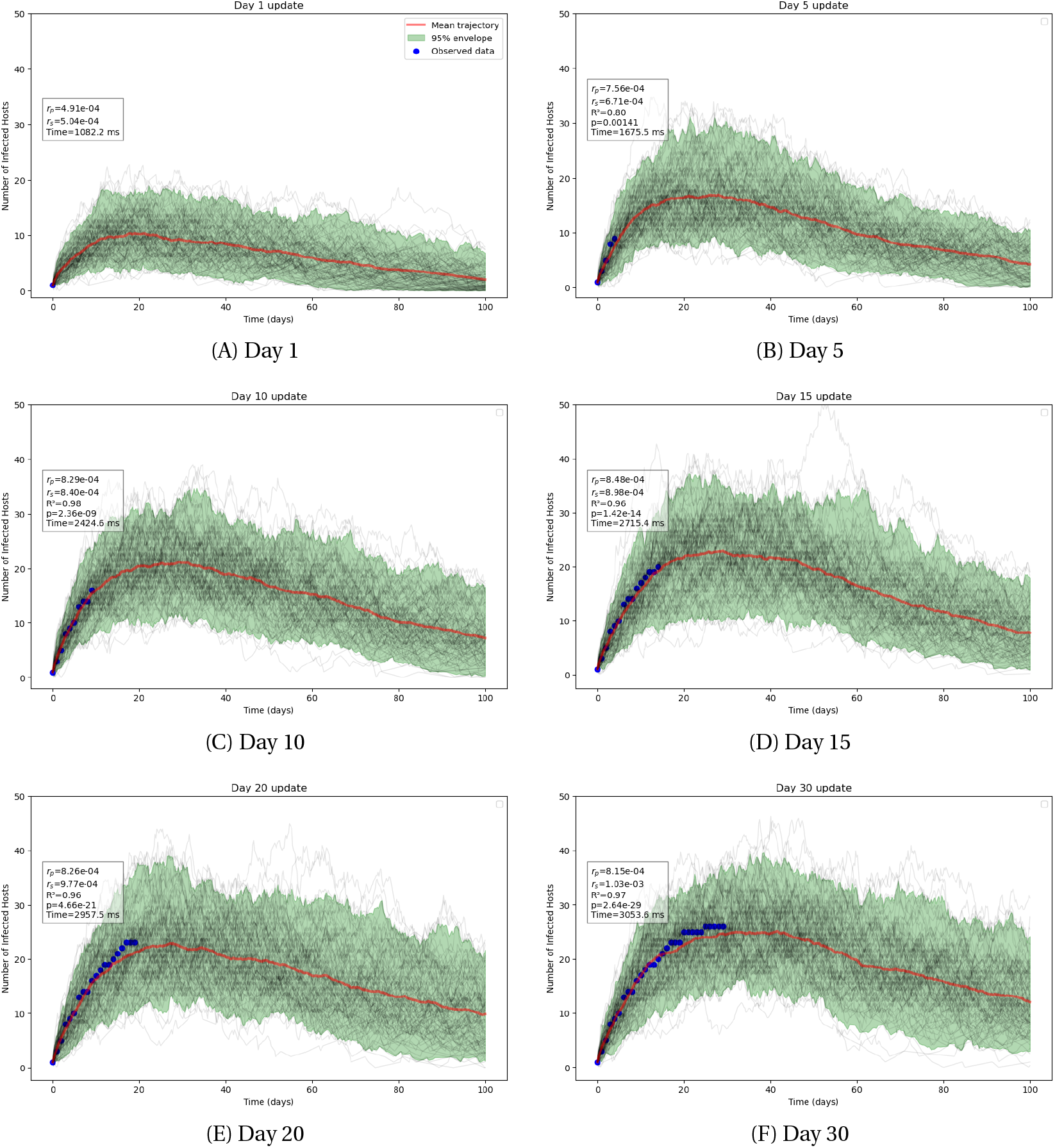
Real-time, day-by-day predictive ensembles under the posterior-mean parameters: 100 Gillespie simulations (black), ensemblemean trajectory (red), 95% credible envelope (green), and observed counts (blue). Panels: (A) Day 1; (B) Day 5; (C) Day 10; (D) Day 15; (E) Day 20; (F) Day 30. In each panel, the white text box displays 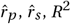, Pearson *p*-value, and computation time (ms).

### 3.2 Cumulative goodness-of-fit

Figure 2 shows the daily *R*^2^ values, accompanied by a 2-day Gaussian-smoothed trend (red). Key observations include:

- On **Day 2**, *R*^2^ *=* 0.78, indicating that two noisy observations do not yet constrain the trajectory fully.
- By **Day 3**, *R*^2^ increases to 0.87, but on **Day 4** it dips to 0.76, reflecting early variability in the data.
- On **Day 5**, *R*^2^ *=* 0.80, and then on **Days 6-7**, it rises to 0.89, showing improving fit once more observations are assimilated.
- From **Day 8** (*R*^2^ *=* 0.88) to **Day 10** (*R*^2^ *=* 0.98), the filter rapidly gains accuracy; Day 9 already reaches 0.90.
- On **Days 11-15**, *R*^2^ remains very high (0.97 on Day 11, 0.92 on Day 12, then 0.96-0.98 through Day 15), indicating that the mean forecast captures over 92% of the variance in those first 15 days.
- From **Day 16** onward, *R*^2^ stays above 0.94, with occasional slight dips (e.g., Day 19: 0.95, Day 23: 0.94), but peaks at 0.99 on Day 24.
- On **Day 29**, *R*^2^ briefly drops to 0.88, likely due to the stochastic nature of the model and the consequent noisy observations, before rebounding to 0.97 on Day 30 and 0.96 on Day 31.
- By **Day 30**, *R*^2^ *=* 0.97, confirming that the ensemble-mean forecast explains nearly all of the variation in the 30 observed points.

### 3.3 Parameter Convergence

Figure 3 depicts both the posterior-mean trajectories and their 95% credible envelopes for *r*_*p*_ (blue curve with light-blue shading) and *r*_*s*_ (green curve with light-green shading) across the 31-day surveillance window. Key features are as follows:

- **Initial phase (Days 1-3):** On Day 1, 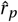 and 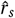 both start near 5.0*×*10^*™*4^. Their envelopes are extremely wide (approximately *±*3 *×* 10^*™*4^) reflecting the uniform prior and minimal data. By Day 3, 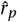 has risen to about 6.5 *×* 10^*™*4^, while 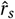 climbs more steeply to about 5.3 *×* 10^*™*4^. Although the means have begun to separate, the overlapping envelopes (Day 3 width *≈ ±*3 *×* 10^*™*4^) indicate substantial uncertainty.
- **Rapid rise (Days 4-9):** Between Days 4 and 9, both parameters ascend quickly. By Day 7, 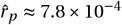 and also 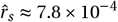, with their envelopes narrowing to roughly *±*1.5 *×* 10^*™*4^. Crucially, *r*_*s*_ overtakes *r*_*p*_ around Day 8, and from Day 9 onward, 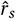 remains above 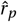. The envelopes have shrunk to *±*1.2 *×* 10^*™*4^, indicating the filter is already focusing on a narrower posterior region.
- **Mid-window stabilization (Days 10-15):** From Day 10 to Day 15, 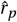 plateaus around 7.8 to 8.3*×*10^*™*4^ while 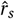 climbs from 8.8 *×*10^*™*4^ to 9.1 *×*10^*™*4^. By Day 12, the 95% envelope for *r*_*p*_ has contracted to approximately *±*1.0 *×* 10^*™*4^ and remains stable-evidence that *r*_*p*_ is becoming well-identified. Meanwhile, *r*_*s*_’s envelope narrows to about *±*1.5 *×* 10^*™*4^ by Day 12 but continues to contract more slowly.
- **Late-window trends (Days 16-31):** After Day 15, 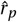 exhibits a gentle decline, falling from *≈* 8.3 *×* 10^*™*4^ on Day 15 to *≈* 8.1*×*10^*™*4^ by Day 31. Its envelope remains tight (about *±*0.8*×*10^*™*4^), indicating high confidence despite the slight downward drift-likely caused by noisy mid-window data points. In contrast, 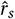 rises almost linearly from *≈* 9.1 *×* 10^*™*4^ on Day 15 to *≈* 10.4 *×* 10^*™*4^ by Day 31. Its envelope narrows to about *±*1.0 *×* 10^*™*4^ by Day 20 and remains roughly that width through Day 31, confirming robust identification of stronger secondary transmission.
- **Relative uncertainty and separation:** Throughout the entire window, *r*_*s*_ maintains a wider 95% envelope than *r*_*p*_. The gap between the blue and green posterior means widens from Day 8 onward, indicating that the filter places increasing weight on secondary transmission (*r*_*s*_) relative to primary transmission (*r*_*p*_). By Day 31, 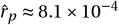 and 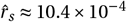.

In summary, both *r*_*p*_ and *r*_*s*_ converge rapidly within the first 10-15 days. The 95% envelopes shrink by more than half over that period, and the posterior means stabilize thereafter. Notably, *r*_*s*_ surpasses *r*_*p*_ on Day 8 and continues to increase, while *r*_*p*_ levels off and then slightly decreases-reflecting the influence of later noisy observations on *r*_*p*_’s posterior, whereas *r*_*s*_ is consistently driven upward toward the true generating value.

### 3.4 Computational Performance

Table 3 reports the wall-clock time for each day’s update on an Intel i7-1185G7 laptop (4 physical/8 logical cores, 31.7 GB RAM, Windows 11). Updates on Days 1-5 run in roughly 1.05-1.90 s, increasing to 2.52 s by Day 10. As more data are assimilated, execution time grows steadily: Days 16-20 take 2.72-2.96 s each, and Days 27-30 require 3.03-3.13 s. Even at maximum data volume, each update completes in under 3.2 s, confirming that the method remains practical for real-time epidemic surveillance.

**Table 3.** Execution time (ms) for each day’s particle-filter update on an Intel i7-1185G7 laptop (Windows 11, 31.7 GB RAM). Early updates (Days 1-5) range from about 1.05-1.90 s; by Day 10 they reach approximately 2.52 s. As the data window grows, Days 16-20 cluster around 2.72-2.96 s, and the last days (Days 27-30) require about 3.03-3.13 s. Even at peak load, each update remains under 3.2 s, demonstrating that the memory-based filter scales efficiently for real-time tracking.

| Day | Time (ms) | Day | Time (ms) | Day | Time (ms) | Day | Time (ms) |
| --- | --- | --- | --- | --- | --- | --- | --- |
| 1 | 1058.2 | 9 | 2424.6 | 17 | 2802.6 | 25 | 3111.2 |
| 2 | 1158.3 | 10 | 2522.9 | 18 | 2765.1 | 26 | 3053.4 |
| 3 | 1360.8 | 11 | 2550.5 | 19 | 2957.5 | 27 | 3037.9 |
| 4 | 1675.5 | 12 | 2786.8 | 20 | 2920.8 | 28 | 3135.2 |
| 5 | 1898.9 | 13 | 2909.5 | 21 | 2940.5 | 29 | 3053.6 |
| 6 | 2102.5 | 14 | 2715.4 | 22 | 2971.2 | 30 | 3059.5 |
| 7 | 2230.5 | 15 | 2712.4 | 23 | 3043.1 | 31 | — |
| 8 | 2368.8 | 16 | 2725.9 | 24 | 3047.7 |  |  |

### 3.5 Summary of Findings

The memory-based particle filter demonstrated rapid learning, robust forecasting, and efficient computational performance when applied to real-time epidemic tracking. Key findings include:

1. **Forecast accuracy:** Real-time forecasts rapidly improved with sequential data assimilation. By Day 5,all observations fell within the 95% credible envelope (Fig. 1A-F). From Day 10 onward, ensemble-mean predictions closely tracked observations (within *±*1 host), with the *R*^2^ coefficient consistently exceeding 0.94 (Fig. 2). This confirms the filter’s ability to produce well-calibrated, high-fidelity forecasts.

**Figure 2.**
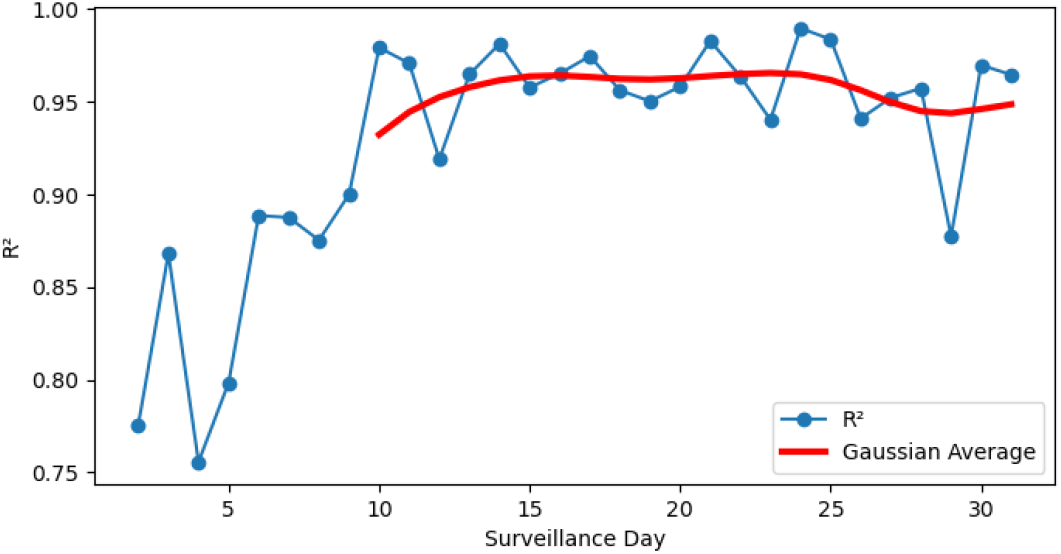
Temporal evolution of the coefficient of determination *R*^2^ comparing the ensemble-mean prediction to observed data up to each day. The red curve is a 2-day Gaussian smoothing. From Day 5 onward, *R*^2^ *≥* 0.88; by Day 9, *R*^2^ *>* 0.95; and from Day 10-30, *R*^2^ *≈* 0.94-0.99.

**Parameter convergence:** Posterior estimates for transmission rates (*r*_*p*_, *r*_*s*_) converged within 10-15 days (Fig. 3). The 95% credible intervals narrowed by more than 50% during this period, with *r*_*s*_ (secondary transmission) surpassing *r*_*p*_ (primary transmission) by Day 8. Final estimates 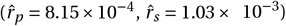 approached the true generating values (8.5 *×* 10^*™*4^, 1.1 *×* 10^*™*3^).

**Figure 3.**
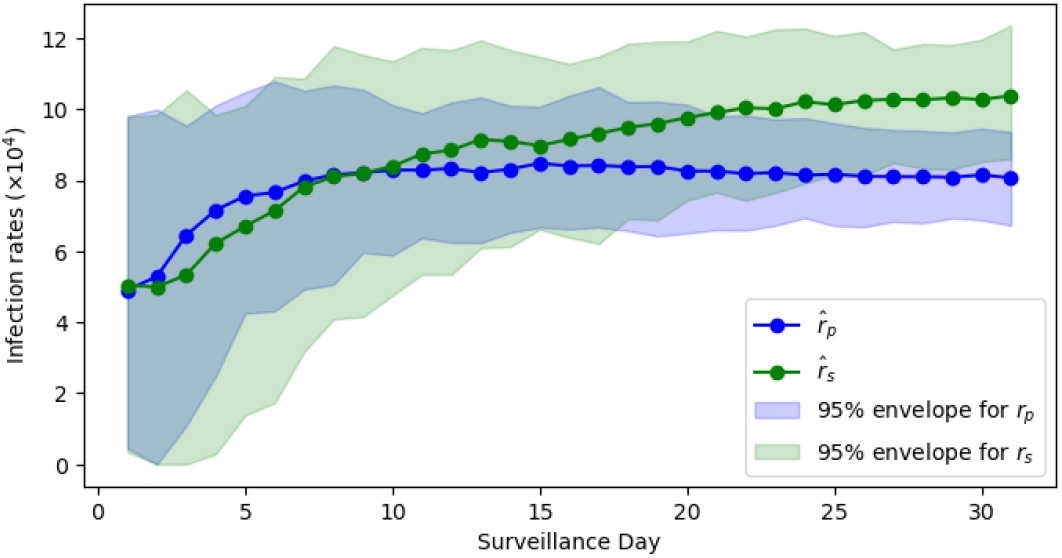
Posterior-mean estimates of *r*_*p*_ (blue) and *r*_*s*_ (green), each scaled by 10^4^, over Days 131. Shaded regions denote the 95% credible envelope for each parameter.

**Goodness-of-fit:** The ensemble-mean trajectory explained nearly all variance in the data (*R*^2^ *≥* 0.94) after Day 10 (Fig. 2). Transient dips in *R*^2^ (e.g., Day 29: 0.88) reflected stochastic noise but recovered swiftly, underscoring the filter’s resilience.

**Computational efficiency:** Daily updates executed in 1.05-3.13 seconds (Table 3), scaling linearly with data accumulation. Even at peak load (Day 31: 3.06 s), performance remained practical for real-time deployment on standard hardware.

Collectively, these results validate the filter’s capability to deliver accurate, uncertainty-quantified forecasts with rapid parameter identification, enabling reliable real-time epidemic surveillance.

## 4 Discussion and conclusion

The sequential Monte Carlo method introduced here demonstrates that real-time, day-by-day parameter estimation in a stochastic plant epidemic model can rapidly converge to near-true values, even when counts are low. By approximately Day 10-15, posterior means for both primary (*r*_*p*_) and secondary (*r*_*s*_) infection rates were within respectively *∼* 1% and *∼* 18% of their ground-truth values, and both were within *∼* 6% by day 31. This rapid convergence was made possible by memory-based sampling within bootstrap filters, which focuses computational effort on high-posterior-density regions without requiring exhaustive resampling at every time-step. In contrast to retrospective, batch-based calibration methods, such as least-squares fitting to full incidence-curve ODE models, my approach dynamically refines parameter estimates as each new observation arrives, thereby minimizing lag between data collection and model adjustment.

Beyond parameter recovery, high predictive accuracy was achieved very early in the epidemic trajectory. Even with a small number of discrete counts, reliable short-term forecasts emerged and uncertainty was explicitly represented via stochastic simulation. This stands in contrast to deterministic ODE-based forecasts, which often underestimate variability in early epidemic stages.

From a computational stand-point, daily updates required between roughly 1s (Day 1) and 3s (Day 31) on a standard laptop, making the method feasible for real-time decision support in field settings. Because the epidemic model is relatively parsimonious (two reaction channels) and memory-based sampling concentrates particles near high-likelihood regions, scalability remains acceptable even as the observational window widens. In operational terms, this suggests that robust parameter inference and forecasting could be embedded within automated surveillance platforms, thereby enabling adaptive management of plant diseases.

Nevertheless, several limitations warrant consideration. First, my validation used synthetic data generated from nearly the same mean-field ODE used to prescribe daily change, so inference performance may be optimistic relative to real-world scenarios where model misspecification and observation biases occur [24]. Second, the simplified epidemic model omits latent (exposed) compartments, spatial structure, and host demography features known to affect transmission dynamics in many pathosystems. Although latent-period effects can sometimes be approximated via adjustments to the susceptible pool or infection rates, explicit SEI formulations may be necessary for diseases with prolonged incubation. Similarly, spatial heterogeneity can skew parameter estimates if local clustering of infection is pronounced.

Future work should extend the current framework to accommodate field data with irregular sampling intervals, multiple plot replicates, and covariate information (e.g., weather or cultivar susceptibility). Integration of geospatially explicit models, perhaps via spatially constrained particle filters, could capture local inoculum sources and dispersal pathways without resorting to fully spatially explicit PDEs. Furthermore, hybrid inference schemes that employ fast ODE-based warm starts followed by stochastic refinement may accelerate early updates when infection counts remain low. Finally, empirical validation against archival surveillance datasets will be essential to assess performance under real-world noise, reporting delays, and control interventions.

In conclusion, I have presented a sequential-Bayesian approach with memory for estimating key transmission parameters in a stochastic plant disease model, achieving rapid convergence and accurate short-term forecasts while explicitly quantifying uncertainty. By combining the Gillespie stochastic algorithm with memory-based particle resampling, my method overcomes limitations of retrospective, batch-based calibration and offers a scalable solution for real-time epidemiological decision support. With further extensions to account for latent periods, spatial structure, and heterogeneous data streams, sequential Monte Carlo approaches have the potential to enhance parameter inference and forecasting in plant-health surveillance.

## Data and Code availability

All simulation code, synthetic data sets and analysis scripts are archived on GitHub: https://github.com/israeltankam/Memory-based-incremental-parameter-update-of-a-generic-stochastic-plant-epidemic-model.

Readers can reproduce all figures and run the sequential Monte Carlo framework on their own data.

## Acknowledgments

I gratefully acknowledge the use of OpenAI’s ChatGPT (gpt-4 model) in the preparation of this manuscript. ChatGPT was employed to proofread text for typos and clarity, to help debug and refine code, and to generate prototype code snippets when needed. All AI-generated suggestions were reviewed, edited, and approved before inclusion.

## A Gillespie formulation of the generalized plant disease model

I consider the stochastic simulation of

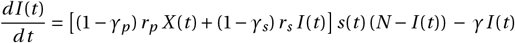

via the Gillespie algorithm.

### 1. State and Events

- *I* (*t*): current number of infected hosts;
- *N* : total (constant) host population;
- *X* (*t*): external inoculum (continuous);
- *s*(*t*): time-varying susceptibility modifier.

I define two reaction events:

| Reaction | Description | $\Delta I$ |
| --- | --- | --- |
| 1. Infection | primary + secondary infection | +1 |
| 2. Removal | removal / control of an infected host | -1 |

### 2. Propensities

At time *t*, with state *I* :

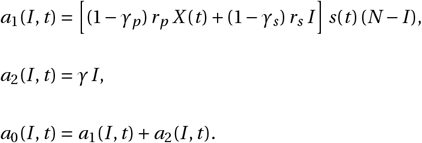

### 3. Gillespie Algorithm

1. Initialize: *t ← t*_0_, *I ← I*_0_.
2. Compute propensities *a*_1_, *a*_2_, *a*_0_.
3. Draw *u*_1_, *u*_2_ *∼* Uniform(0, 1).
4. Compute time to next event:

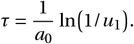
5. Determine which reaction occurs: choose *j ∈* {1, 2} such that

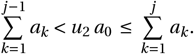
6. Update:

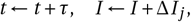

where Δ*I*_1_ *= +*1, Δ*I*_2_ *= ™*1.
7. Update time-dependent inputs *X* (*t*), *s*(*t*) as needed.
8. Repeat until final time or *I =* 0.

## B Daily update figures

Below are all of the daily-update plots, arranged so that they span multiple pages. Each panel shows 100 Gillespie trajectories (black), the ensemble-mean trajectory (red), the 95 % predictive envelope (green), and the observed infected counts up to that day (blue). The white text box in each subplot displays 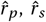, *R*^2^, Pearson *p*-value, and computation time (ms).

**Figure 4.**
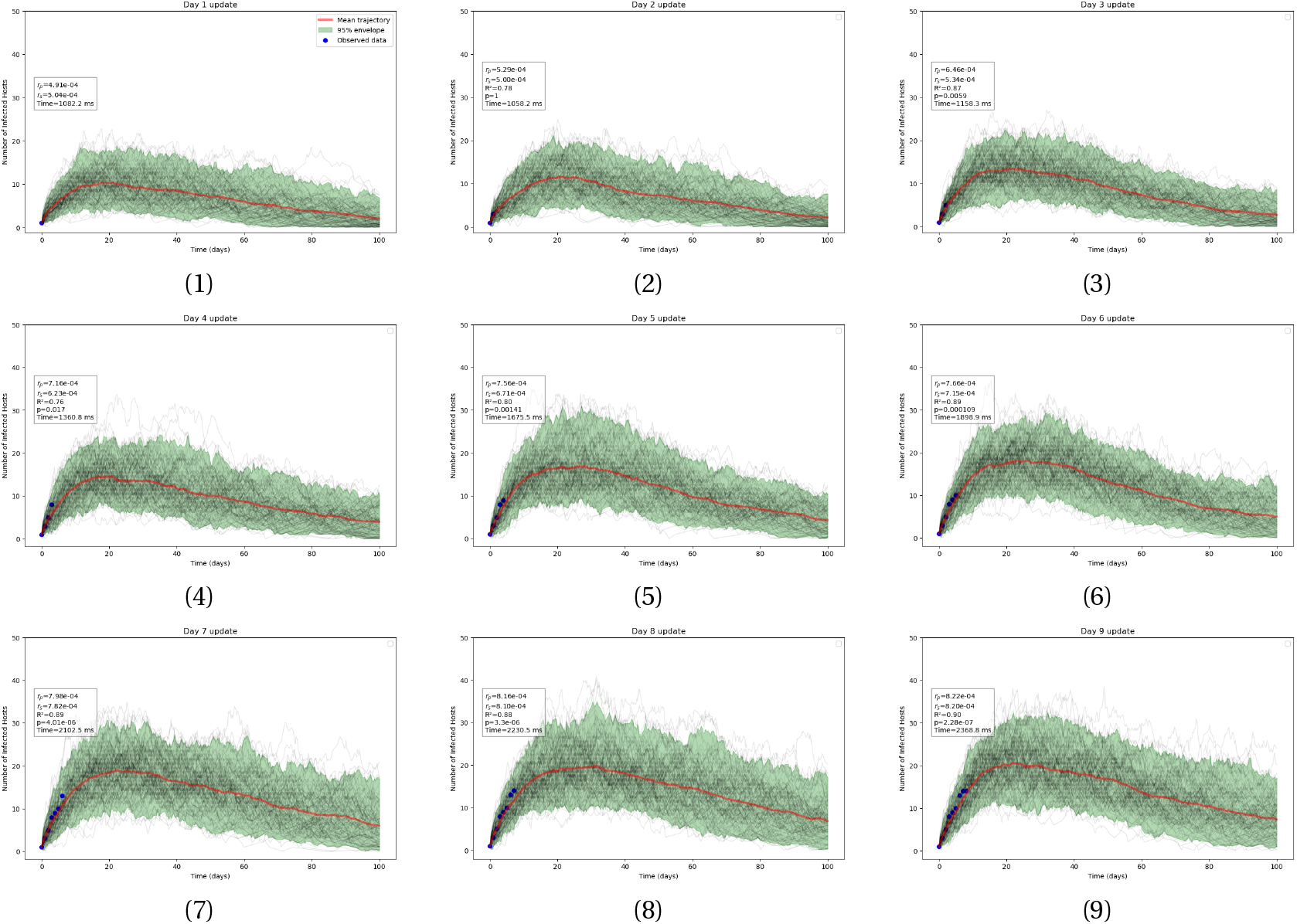
Daily update figures for Days 1-9. Each panel displays: 100 Gillespie simulations (black), the ensemble-mean trajectory (red), the 95 % envelope (green), and observed counts (blue). In each subplot, the white text box shows 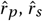, *R*^2^, Pearson *p*-value, and computation time (ms).

**Figure 5.**
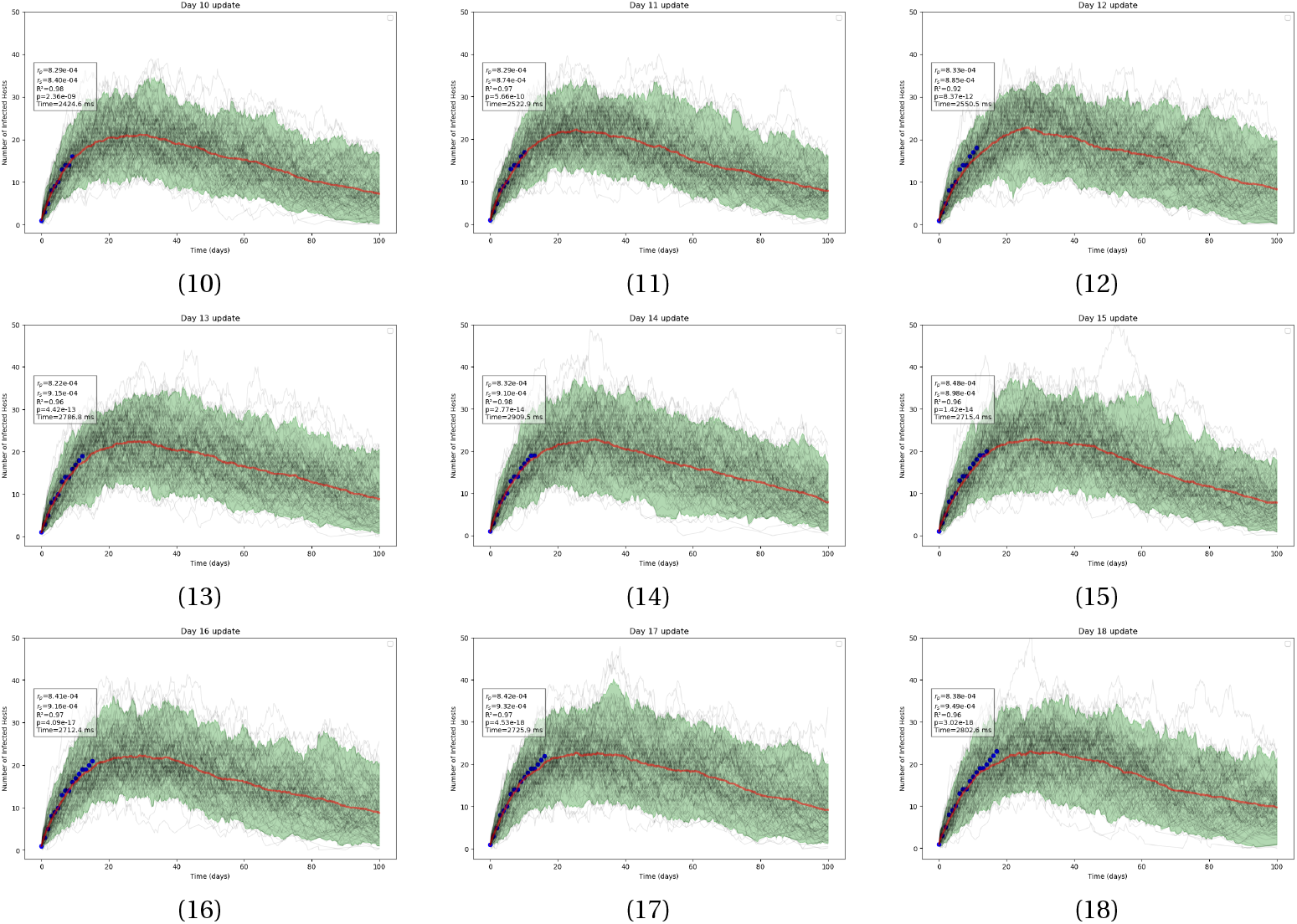
Daily update figures for Days 10-18. Each panel displays: 100 Gillespie simulations (black), the ensemble-mean trajectory (red), the 95 % envelope (green), and observed counts (blue). In each subplot, the white text box shows 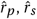, *R*^2^, Pearson *p*-value, and computation time (ms).

**Figure 6.**
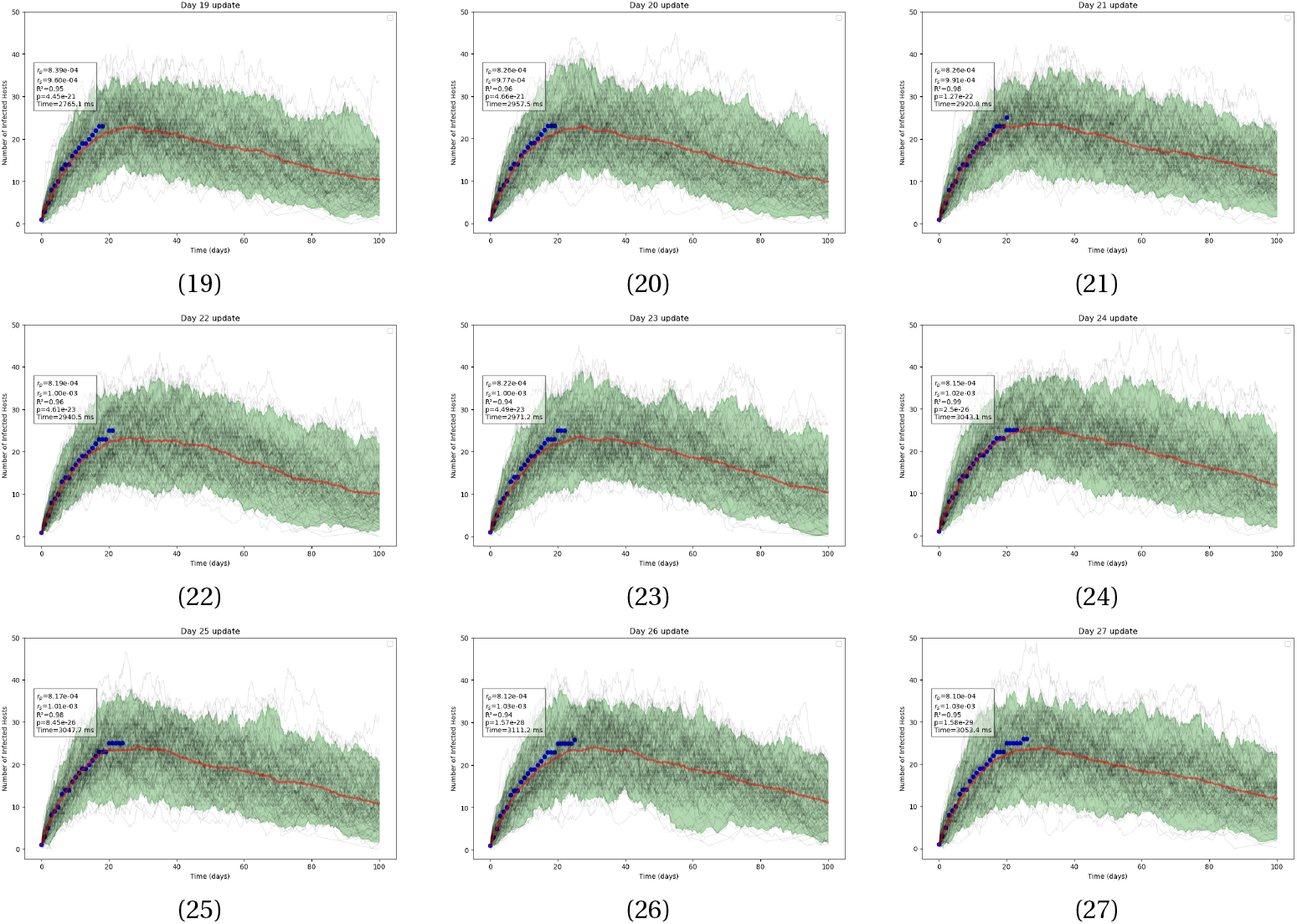
Daily update figures for Days 19-27. Each panel displays: 100 Gillespie simulations (black), the ensemble-mean trajectory (red), the 95 % envelope (green), and observed counts (blue). In each subplot, the white text box shows 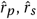, *R*^2^, Pearson *p*-value, and computation time (ms).

**Figure 7.**
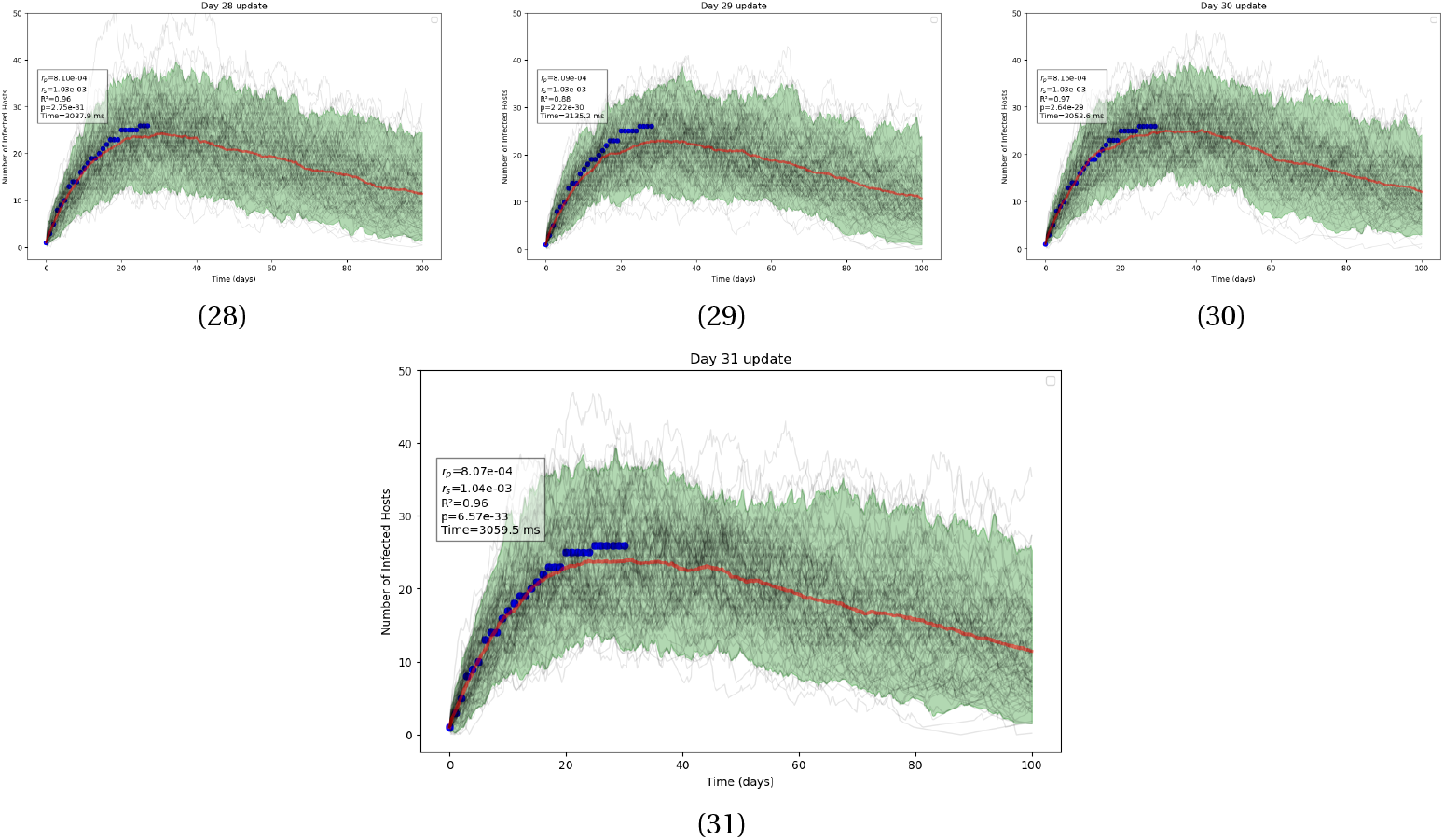
Daily update figures for Days 28-31. Each panel displays: 100 Gillespie simulations (black), the ensemble-mean trajectory (red), the 95 % envelope (green), and observed counts (blue). In each subplot, the white text box shows 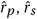, *R*^2^, Pearson *p*-value, and computation time (ms).

## Notes

https://github.com/israeltankam/Memory-based-incremental-parameter-update-of-a-generic-stochastic-plant-epidemic-model

### Competing Interest Statement

The authors have declared no competing interest.

## References

[1] Alfred B. Amendolara, David Sant, Horacio G. Rotstein, and Eric Fortune. Lstm-based recurrent neural network provides effective short term flu forecasting. BMC Public Health, 23(1), September 2023. ISSN 1471-2458. doi: 10.1186/s12889-023-16720-6. URL http://dx.doi.org/10.1186/s12889-023-16720-6.

[2] D Anderson and K Burnham. Model selection and multi-model inference. Second. NY: Springer-Verlag, 63 (2020):10, 2004.

[3] M.S. Arulampalam, S. Maskell, N. Gordon, and T. Clapp. A tutorial on particle filters for online nonlinear/non-gaussian bayesian tracking. IEEE Transactions on Signal Processing, 50(2):174–188, 2002. ISSN 1053-587X. doi: 10.1109/78.978374. URL http://dx.doi.org/10.1109/78.978374.

[4] H. J. Aust and J. Kranz. Experiments and Procedures in Epidemiological Field Studies, page 7–17. Springer Berlin Heidelberg, 1988. ISBN 9783642955341. doi: 10.1007/978-3-642-95534-1_2. URL http://dx.doi.org/10.1007/978-3-642-95534-1_2.

[5] D. J. Bailey and C. A. Gilligan. Dynamics of primary and secondary infection in take-all epidemics. Phytopathology®, 89(1):84–91, January 1999. ISSN 1943-7684. doi: 10.1094/phyto.1999.89.1.84. URL http://dx.doi.org/10.1094/PHYTO.1999.89.1.84.

[6] Nik J. Cunniffe and Christopher A. Gilligan. Invasion, persistence and control in epidemic models for plant pathogens: the effect of host demography. Journal of The Royal Society Interface, 7(44):439–451, July 2009. ISSN 1742-5662. doi: 10.1098/rsif.2009.0226. URL http://dx.doi.org/10.1098/rsif.2009.0226.

[7] Ruairí Donnelly and Christopher A. Gilligan. A new method for the analysis of access period experiments, illustrated with whitefly-borne cassava mosaic begomovirus. PLOS Computational Biology, 19(8):e1011291, 2023. doi: 10.1371/journal.pcbi.1011291.

[8] Ruairí Donnelly, Israël Tankam, and Chris Gilligan. Plant pathogen profiling with the epipv package. EcoEvoRxiv, 2025. URL https://ecoevorxiv.org/repository/view/9081/.

[9] Kaique dos S Alves, Willian B Moraes, Wellington B da Silva, and Emerson M Del Ponte. Estimation of a time-varying apparent infection rate from plant disease progress curves: A particle filter approach. May 2019. doi: 10.1101/625822. URL http://dx.doi.org/10.1101/625822.

[10] Arnaud Doucet, Nando De Freitas, Neil James Gordon, et al. Sequential Monte Carlo methods in practice, volume 1. Springer, 2001.

[11] Torsten A. Enßlin, Markus Frommert, and Francisco S. Kitaura. Information field theory for cosmological perturbation reconstruction and nonlinear signal analysis. Physical Review D, 80(10):105005, 2009. doi: 10.1103/PhysRevD.80.105005. URL https://journals.aps.org/prd/abstract/10.1103/PhysRevD.80.105005.

[12] Wilfried Gabriel and Reinhard Bürger. Survival of small populations under demographic stochasticity. Theoretical Population Biology, 41(1):44–71, 1992. doi: 10.1016/0040-5809(92)90049-Y.

[13] G J Gibson, C A Gilligan, and A Kleczkowski. Predicting variability in biological control of a plant—pathogen system using stochastic models. Proceedings of the Royal Society of London. Series B: Biological Sciences, 266 (1430):1743–1753, 1999. doi: 10.1098/rspb.1999.0841.

[14] Daniel T. Gillespie. Exact stochastic simulation of coupled chemical reactions. The Journal of Physical Chemistry, 81(25):2340–2361, December 1977. ISSN 1541-5740. doi: 10.1021/j100540a008. URL http://dx.doi.org/10.1021/j100540a008.

[15] William R Gillespie. Noncompartmental versus compartmental modelling in clinical pharmacokinetics. Clinical pharmacokinetics, 20(4):253–262, 1991. doi: 10.2165/00003088-199120040-00001.

[16] C.A. Gilligan. Mathematical Modelling of Crop Disease. Advances in plant pathology. Academic Press, 1985. ISBN 9780120337033.

[17] Tilmann Gneiting and Adrian E Raftery. Strictly proper scoring rules, prediction, and estimation. Journal of the American Statistical Association, 102(477):359–378, March 2007. ISSN 1537-274X. doi: 10.1198/016214506000001437. URL http://dx.doi.org/10.1198/016214506000001437.

[18] N.J. Gordon, D.J. Salmond, and A.F.M. Smith. Novel approach to nonlinear/non-gaussian bayesian state estimation. IEE Proceedings F Radar and Signal Processing, 140(2):107, 1993. ISSN 0956-375X. doi: 10.1049/ip-f-2.1993.0015. URL http://dx.doi.org/10.1049/ip-f-2.1993.0015.

[19] Trevor Hastie, Robert Tibshirani, Jerome H Friedman, and Jerome H Friedman. The elements of statistical learning: data mining, inference, and prediction, volume 2. Springer, 2009.

[20] Skylar R. Hopkins, Arietta E. Fleming-Davies, Lisa K. Belden, and Jeremy M. Wojdak. Systematic review of modelling assumptions and empirical evidence: Does parasite transmission increase nonlinearly with host density? Methods in Ecology and Evolution, 11(4):476–486, February 2020. ISSN 2041-210X. doi: 10.1111/2041-210x.13361. URL http://dx.doi.org/10.1111/2041-210X.13361.

[21] Genshiro Kitagawa. Monte carlo filter and smoother for non-gaussian nonlinear state space models. Journal of Computational and Graphical Statistics, 5(1):1–25, March 1996. ISSN 1537-2715. doi: 10.1080/10618600.1996.10474692. URL http://dx.doi.org/10.1080/10618600.1996.10474692.

[22] A Kleczkowski, D J Bailey, and Christopher A Gilligan. Dynamically generated variability in plant-pathogen systems with biological control. Proceedings of the Royal Society of London. Series B: Biological Sciences, 263 (1371):777–783, 1996.

[23] Joseph Lee Rodgers and W. Alan Nicewander. Thirteen ways to look at the correlation coefficient. The American Statistician, 42(1):59–66, February 1988. ISSN 1537-2731. doi: 10.1080/00031305.1988.10475524. URL http://dx.doi.org/10.1080/00031305.1988.10475524.

[24] Daniel E Platt, Laxmi Parida, and Pierre Zalloua. Lies, gosh darn lies, and not enough good statistics: why epidemic model parameter estimation fails. Scientific Reports, 11(1):408, 2021. doi: 10.1038/s41598-020-79745-6.

[25] Rachel Russell and Nik J. Cunniffe. Optimal control prevents itself from eradicating stochastic disease epidemics. PLOS Computational Biology, 21(2):e1012781, 2025. doi: 10.1371/journal.pcbi.1012781.

[26] Israël Tankam-Chedjou, Frédéric Grognard, Jean Jules Tewa, and Suzanne Touzeau. Modelling and control of a banana soilborne pest in a multi-seasonal framework. Mathematical Biosciences, 322:108324, 2020. doi: 10.1016/j.mbs.2020.108324.

[27] Robin N. Thompson, Christopher A. Gilligan, and Nik J. Cunniffe. Control fast or control smart: When should invading pathogens be controlled? PLOS Computational Biology, 14(2):e1006014, 2018. doi: 10.1371/journal.pcbi.1006014.

[28] J. E. Truscott, C. R. Webb, and C. A. Gilligan. Asymptotic analysis of an epidemic model with primary and secondary infection. Bulletin of Mathematical Biology, 59(6):1101–1123, November 1997. ISSN 1522-9602. doi: 10.1007/bf02460103. URL http://dx.doi.org/10.1007/BF02460103.

[29] Minghui Wang and Tong Li. Pest and disease prediction and management for sugarcane using a hybrid autoregressive integrated moving average—a long short-term memory model. Agriculture, 15(5):500, February 2025. ISSN 2077-0472. doi: 10.3390/agriculture15050500. URL http://dx.doi.org/10.3390/agriculture15050500.

[30] C J Willmott and K Matsuura. Advantages of the mean absolute error (mae) over the root mean square error (rmse) in assessing average model performance. Climate Research, 30:79–82, 2005. ISSN 1616-1572. doi: 10.3354/cr030079. URL http://dx.doi.org/10.3354/cr030079.

